# The Impact of Pf Bacteriophages on the Fitness of *Pseudomonas aeruginosa*; A Mathematical Modeling Approach

**DOI:** 10.1101/2020.08.30.272203

**Authors:** Julie D. Pourtois, Michael J. Kratochvil, Qingquan Chen, Naomi L. Haddock, Elizabeth B. Burgener, Giulio A. De Leo, Paul L. Bollyky

**Author notes:** **Corresponding author:** Julie Pourtois.

## Abstract

*Pseudomonas aeruginosa* (*Pa*) is a major bacterial pathogen responsible for chronic lung infections in cystic fibrosis patients. Recent work by ourselves and others has implicated Pf bacteriophages, non-lytic filamentous viruses produced by *Pa*, in the chronicity and severity of *Pa* infections. Pf phages act as structural elements in *Pa* biofilms and sequester aerosolized antibiotics, thereby contributing to antibiotic tolerance. Consistent with a selective advantage in this setting, the prevalence of Pf+ bacteria increases over time in these patients. However, the production of Pf phages comes at a metabolic cost to bacteria, such that Pf+ strains grow more slowly than Pf- strains in vitro. Here, we use a mathematical model to investigate how these competing pressures might influence the relative abundance of Pf+ versus Pf- strains in different settings. Our model predicts that Pf+ strains of *Pa* can only outcompete Pf- strains if the benefits of phage production falls solely onto Pf+ strains and not onto the overall bacterial community in the lung. Further, phage production only leads to a net positive gain in fitness at antibiotic concentrations slightly above the minimum inhibitory concentration (i.e., concentrations for which the benefits of antibiotic sequestration outweigh the metabolic cost of phage production), but which are not lethal for Pf+ strains. As a result, our model predicts that frequent administration of intermediate doses of antibiotics with low decay rates favors Pf+ over Pf- strains. These models inform our understanding of the ecology of Pf phages and suggest potential treatment strategies for Pf+ *Pa* infections.

**Importance:** Filamentous phages are a frontier in bacterial pathogenesis, but the impact of these phages on bacterial fitness is unclear. In particular, Pf phages produced by *Pa* promote antibiotic tolerance but are metabolically expensive to produce, suggesting that competing pressures may influence the prevalence of Pf+ versus Pf- strains of *Pa* in different settings. Our results identify conditions likely to favor Pf+ strains and thus antibiotic tolerance. This study contributes to a better understanding of the unique ecology of filamentous phages and may facilitate improved treatment strategies for combating antibiotic tolerance.

## Introduction

*Pseudomonas aeruginosa* (*Pa*) infections, particularly those that are antibiotic resistant or antibiotic tolerant, are responsible for growing health care expenses and extensive mortality (1). *Pa* infections are particularly problematic in cystic fibrosis (CF), an inherited disease associated with defective ion transport and thick, tenacious airway secretions (2–5). The establishment of a chronic *Pa* infection often occurs early in life and evolves into an entrenched and highly damaging condition in adult CF patients. By adulthood, nearly 60% of CF patients have chronic *Pa* infections (6) with 25% of these infections harboring antibiotic-resistant *Pa* strains(7). The microbial ecology of the lung is an important variable in clinical outcomes in CF patients (8, 9). Understanding the forces that influence the progression of these infections is critical for developing effective clinical treatments. Mathematical models can yield important insights into these dynamics (10–15).

*Pa* is particularly pathogenic because of its ability to form robust biofilms. Biofilms are viscous conglomerates of polymers and microbial communities that allow *Pa* to colonize airways (16). Once *Pa* biofilm infections are established in CF lungs, they are nearly impossible to eradicate (17, 18). Many antibiotics have limited penetration through biofilms (19) such that the bacteria encased within are antibiotic tolerant (i.e., able to survive exposure to antimicrobials) in comparison to planktonic bacteria (20, 21). Over time, this reduction in effective antimicrobial activity favors the development of antibiotic resistance (20, 22–24) (i.e., the ability to proliferate despite the presence of antibiotics (20)). Most CF patients rely on intermittent courses of inhaled, anti-pseudomonal antibiotics, such as tobramycin and aztreonam (25). While there are guidelines to guide therapy, the particular regimen is decided by the provider on a case-by-case basis with mixed clinical success (26–28).

We have identified novel roles for Pf bacteriophages in *Pa* biofilm formation and function (7, 29, 30). Pf phages are filamentous, single stranded DNA viruses in the genus *Inovirus*. Unlike many bacteriophages that lyse their bacterial hosts, Pf phages are produced without lysis (31) and are reported to contribute to *Pa* virulence (32). We have previously reported that Pf phages promote the ordering of biofilm polymers into a liquid crystalline structure that prevents antibiotic diffusion and inhibits bacterial clearance (33–35). In addition, some antibiotics are bound to Pf phage structures and are prevented from killing embedded bacteria (29, 36). This liquid crystalline organization has also been observed in other filamentous phages (37).

Pf phages may contribute to the disease burden associated with *Pa* infections in humans (34, 38). Many (between 36-61%) *Pa* clinical isolates produce Pf phages (7, 39, 40), and the presence of these in the sputum of CF patients is positively correlated with chronicity of *Pa* infections (7, 29) and worse declines in pulmonary function (7, 35, 41). Pf phages likewise enhance the virulence of *Pa* infections in animal models (29), including in airway infections (7).

Consistent with a selective advantage in CF lungs, the prevalence of Pf phages increases with patient age and disease severity. We observed that less than 10% of children with CF who are infected with Pa have Pf+ strains, while over 40% of adults and 100% of 10 pre-lung transplant patients have Pf+ strains (7). The prevalence of Pf phages likewise increases over time in patients with chronic Pa wound infections (42). Together these data point to selective advantages for Pf+ *Pa* strains in the setting of chronic infection.

Of note, in a longitudinal cohort study of CF patients we find that the acquisition of Pf phage represents the appearance of a new strain of *Pa*. In our phylogenetic analyses we have not observed the de novo acquisition of Pf phage by an existing, un-parasitized strain of *Pa* (unpublished results). This suggests that strains of Pa may be highly adapted to carry Pf phages. Further, it suggests that to understand the impact of Pf phages on *Pa* fitness we should consider the competitive interactions of Pf+ versus Pf- *Pa* strains.

The production of Pf phages comes at a steep metabolic cost to the *Pa* strains that produce them. Pf phages are abundantly expressed in Pf+ *Pa* biofilms (43) and CF sputum at an average of 10^7^ copies/mL (33, 35). Pf+ strains of *Pa* consequently grow more slowly due to the energetic demands of phage production (34, 35, 44, 45). The impact of these competing pressures on *Pa* fitness is therefore unclear.

In this study we use a mathematical model to investigate the competitive dynamics of Pf+ and Pf-strains of *Pa*. In particular, we compare the fitness of Pf+ and Pf- strains when exposed to different concentrations of tobramycin in CF lungs and for different metabolic costs. Our data predict that for Pf+ strains to have a selective advantage, antibiotic sequestration is essential and must only benefit the phage-producing bacteria. In addition, our results suggest that Pf production is most advantageous when bacteria are continuously exposed to intermediate concentrations of antibiotics, consistent with the type of environment created by some CF antibiotic treatment regimens.

## Methods

### General framework

We envisioned two alternative (i.e., mutually exclusive) types of interaction between Pf+ and Pf- strains: 1) *direct competition* in a mixed population of Pf- and Pf+ bacteria strains coexisting in the same location (e.g., in direct contact and within the same portion of the lung airway) and 2) no direct interaction, with Pf- and Pf+ strains coexisting in the same patient but established in different (i.e., non-overlapping) locations, e.g., spatially separated infection sites in the lungs (Figure 1) – hereafter referred to also as *indirect competition*. While Pf- and Pf+ do not interact directly, their relative growth rates and abundances affect their probability of transmission to non-infected areas of the lungs.

**Figure 1.**
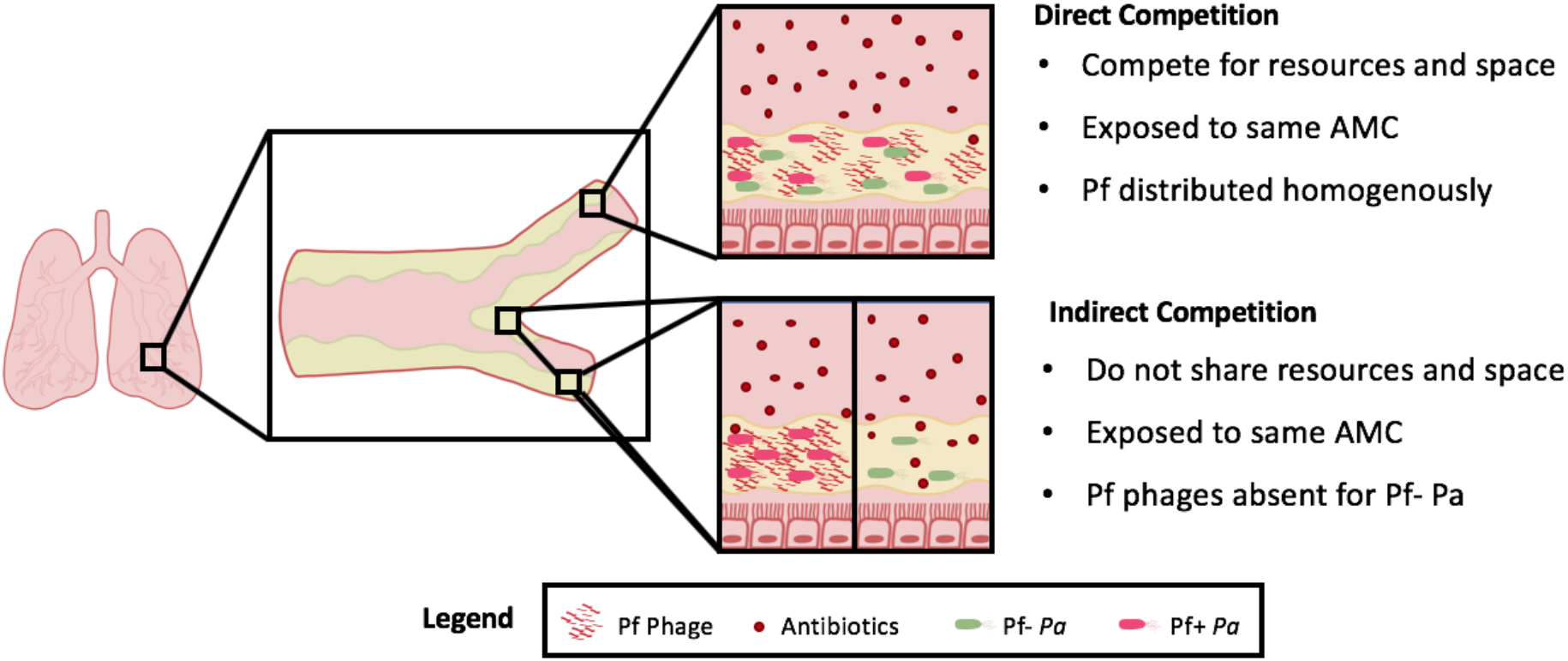
Modeling of the different competitive relationships between Pf+ and Pf- *Pa* bacterial strains. Strains can either compete directly (i.e., when the strains are co-localized), or indirectly (i.e., the strains are not co-localized, but within the same tissue). In both cases, the anti-microbial concentration (AMC) during treatment is the same. However, sequestration of antibiotic by phages leads to a lower concentration of *free* antibiotics capable of inducing bacterial cell death. Our model assumes that the lowering of free antibiotic by sequestration benefits any *Pa* bacterial cells within the location, regardless of whether that cell produces Pf phages.

In direct competition, both Pf- and Pf+ must compete for the same nutrients and space. As filamentous phages are generally highly specific (46, 47), we assumed that the Pf- strain cannot be infected by phages produced by the Pf+ strain. However, under antibiotic treatment, the Pf- strain can benefit from the protective biofilm of filamentous phages produced by the Pf+ strain coexisting in the same location.

In indirect competition, there is no inter-strain competition for nutrients and space and we assumed that, under antibiotic treatment, the Pf- strain does not benefit from the protective effect of phages produced elsewhere by the Pf+ strain.

### Model description

#### Bacterial growth

In the absence of antibiotics, bacteria *B_i_* [CFU/ml] from each strain *i* = Pf+ or Pf-, replicate at a density-dependent rate Φ(*B_tot_*) and die at the per-capita rate *δ_B_*:

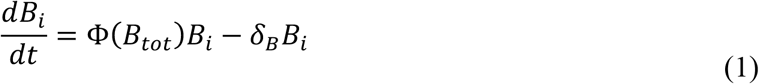

The per-capita replication rate Φ(*B_tot_*) is a decreasing logistic function of the total bacteria concentration in the local biofilm *B_tot_* (48):

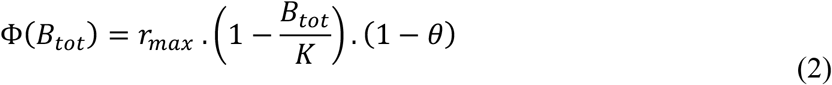

where *r_max_* is the absolute maximum replication rate, K is the bacterial density at which the per capita reproductive rate Φ(*B_tot_*) is equal to zero, and *θ* is the reduction in per-capita replication rate caused by the metabolic cost of phage production in Pf+ bacterial strains only (*θ* is equal to zero for Pf- strains). When both strains are competing directly for space and resources, the total bacteria concentration *B_tot_* is the sum of their concentrations, namely: *B_tot_= B^+^* + *B^-^* When competing indirectly, *B_tot_* is set equal to the local concentration of bacteria, i.e., either the concentration of the Pf+ or of the Pf- strain.

#### Phage production

Filamentous phages *V* [PFU/ml] are produced by Pf+ bacteria at a constant rate *λ* and decay at a rate *δ_V_*:

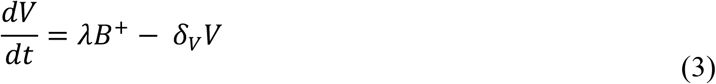

Both *λ* and *δ_V_* were numerically optimized (see **Parametrization**) to obtain viral densities an order of magnitude larger than bacterial densities, consistent with literature reports of *Pa* infections (7).

#### Antibiotic treatment

We consider a range of antibiotic regimens, with different number of doses per day and different antibiotic concentrations per dose. We assume each dose of antibiotics leads to an instantaneous peak *A*_0_ in the sputum. All concentrations discussed in this work represent concentrations in the sputum, which can differ in non-trivial ways from the original concentration administered orally. After administration, antibiotics are removed from the system via both degradation of the drug and clearance due to natural flow through the body. Here we assumed that their concentration follows first order kinetics (49), i.e., it decays at a constant rate *δ_A_* until the next treatment:

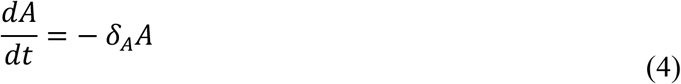

To describe the effect of antibiotics on the bacterial infection, we built upon, and extended, the original model developed by Levin and Udekwu (50). Specifically, the effect of antibiotics in the system is accounted for by modifying equation (1) as follows:

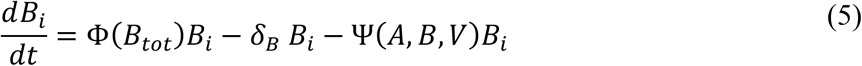

where Ψ(*A*, *B*, *V*) is the per-capita reduction in bacterial growth rate or, equivalently, the increase in mortality rate as a consequence of antibiotics at concentration *A*, bacterial density *B* and phage density *V*–hereafter referred to as antibiotic *killing rate*. Ψ(*A*, *B*, *V*) is here described by a Hill function (51), i.e., an increasing function of the effective antibiotic concentration *A_eff_*(*A*, *V*) that levels off to *ψ*(Φ(*B_tot_*)), the maximum killing rate due to antibiotics:

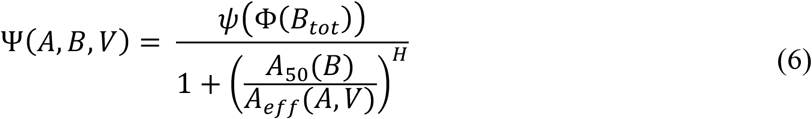

where *A*_50_(*B*) is the antibiotic concentration leading to half *ψ*(Φ(*B_tot_*)), and *H*, the Hill parameter, is proportional to the steepness of the function at *A_eff_*= *A_50_* (50).

The effective killing rate *ψ*(Φ(*B_tot_*)) of the antibiotic is a function of the absolute maximum killing rate Γ and the effect of bacterial replication rate on the antibiotic efficacy (50):

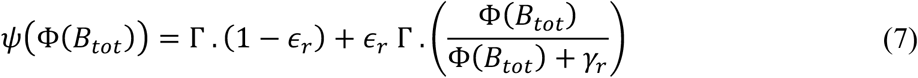

The parameter *ϵ_r_*, bounded between 0-1, describes the degree to which a particular antibiotic’s efficacy depends on the growth rate, i.e., it does not depend upon bacteria reproductive rate Φ(*B_tot_*) when *ϵ_r_* = 0, it depends entirely upon bacteria reproductive rate when *ϵ_r_* = 1. The parameter *γ_r_* represents the growth rate at which the rate of killing is half of its maximum when *ϵ_r_* = 1.

The effective antibiotic concentration is determined by the concentration of antibiotics that are not sequestered by phages. Following Hulme & Trevethick (52), to describe the extent of sequestration, we use the equilibrium expression for ligand-receptor binding as binding dynamics occur at a much faster scale than bacterial killing:

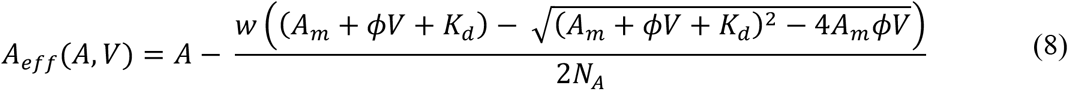

where *ϕ* represents the number of antibiotic binding sites per phage (also referred to as antibiotic sequestration factor), *K_d_* the equilibrium dissociation constant and *A_m_* the antibiotic concentration in molecules per ml. *w* and *N_A_* stand for the molecular weight of the antibiotic and Avogadro’s number, respectively, and are used for the conversion from *μg/ml* to molecule per ml:

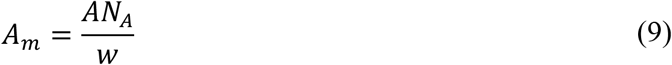

The effective *A*_50_ - the antibiotic concentration leading to half of the maximum killing rate - is here assumed to be an increasing and saturating function of bacterial density *B_tot_* (50), namely:

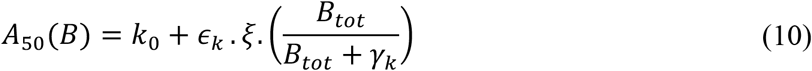

Accordingly, *A*_50_ ranges between *k*_0_ at low bacterial densities and *k*_0_ + *ξ* at high bacterial density, where *ξ* is the maximum additional antibiotic concentration that can be tolerated at high densities, *γ_k_* is the bacterial density at which *A*_50_ increases by half of its maximum amount and *ϵ_k_* is a switch parameter set to zero if *A*_50_ is assumed to be independent from bacterial density, to 1 otherwise.

### Parametrization

The equations in this model are general and can be applied to a variety of environments, bacteria, and antibiotics. Here, we parametrized our model to represent the growing conditions of *Pa* in CF lungs under tobramycin treatment, an antibiotic often used to treat *Pa* infections (Table 1). We chose a value in the middle of the range observed in the literature for most parameters relating to bacterial growth and phage production. For parameters describing the action of antibiotics, we give the range observed across a wide variety of antibiotics, as well as the particular value we used for tobramycin. We performed a global sensitivity analysis to explore how bacterial density responds to variation in parameters around the fixed values used in our model (Figure S1 and S2). In this work, we were concerned with the effect of phage production on fitness and we considered in our analysis a wide range of values for relevant parameters (metabolic cost and antibiotic sequestration constant). Time is measured in hours.

**Table 1.**
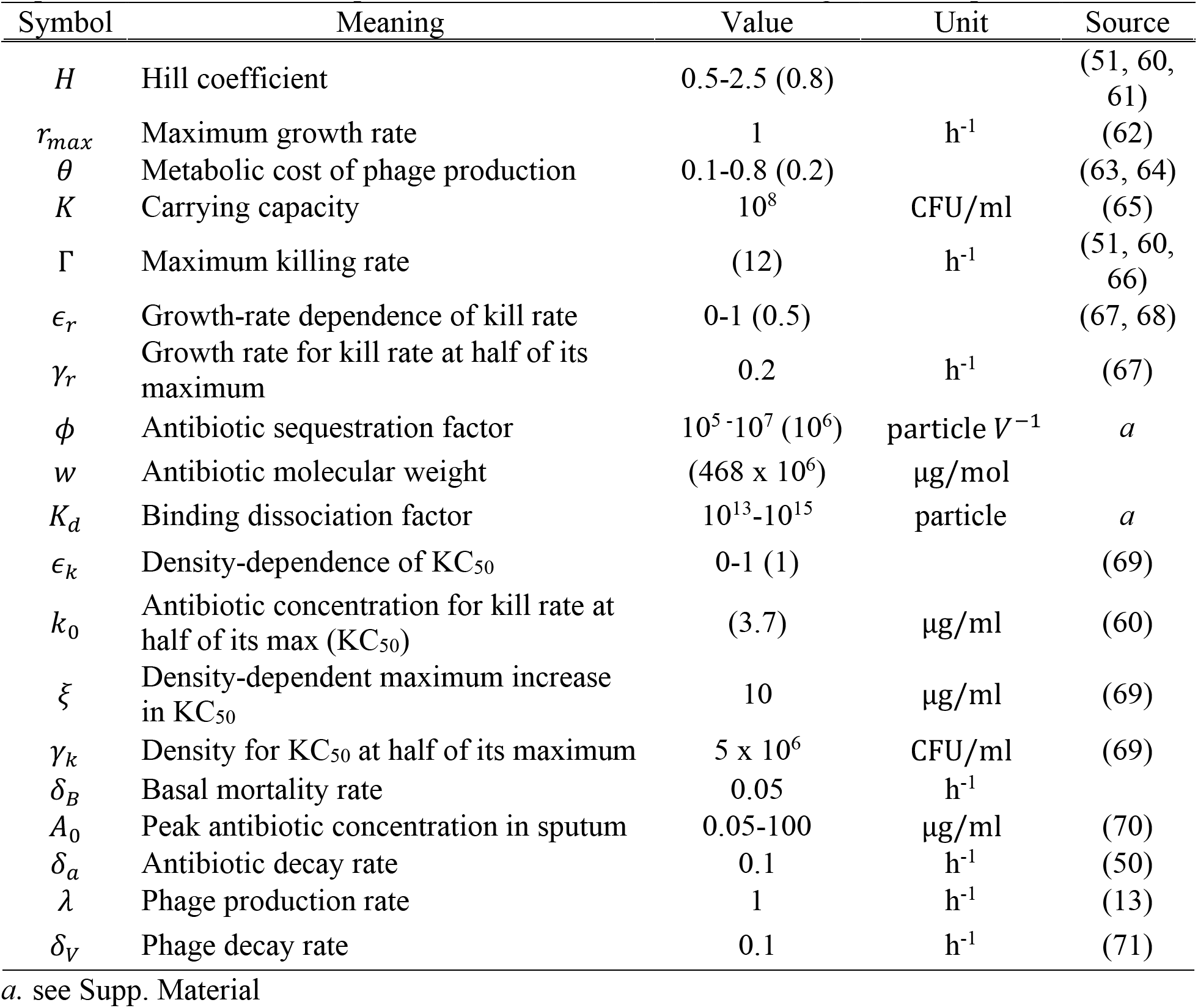
Model parameters. We provide a range of possible values for antibiotic-related parameters and give the exact value we use for tobramycin in parenthesis. We also specify the range of values we consider for the metabolic cost of phage production, the antibiotic sequestration factor, and the peak antibiotic concentration following a dose in sputum.

| Symbol | Meaning | Value | Unit | Source |
| --- | --- | --- | --- | --- |
| $H$ | Hill coefficient | 0.5-2.5 (0.8) | | (51, 60, 61) |
| $r_{max}$ | Maximum growth rate | 1 | $\text{h}^{-1}$ | (62) |
| $\theta$ | Metabolic cost of phage production | 0.1-0.8 (0.2) | | (63, 64) |
| $K$ | Carrying capacity | $10^8$ | CFU/ml | (65) |
| $\Gamma$ | Maximum killing rate | (12) | $\text{h}^{-1}$ | (51, 60, 66) |
| $\epsilon_r$ | Growth-rate dependence of kill rate | 0-1 (0.5) | | (67, 68) |
| $\gamma_r$ | Growth rate for kill rate at half of its maximum | 0.2 | $\text{h}^{-1}$ | (67) |
| $\phi$ | Antibiotic sequestration factor | $10^5$-$10^7$ ($10^6$) | particle $V^{-1}$ | $a$ |
| $w$ | Antibiotic molecular weight | ($468 \times 10^6$) | $\mu\text{g}/\text{mol}$ | |
| $K_d$ | Binding dissociation factor | $10^{13}$-$10^{15}$ | particle | $a$ |
| $\epsilon_k$ | Density-dependence of $\text{KC}_{50}$ | 0-1 (1) | | (69) |
| $k_0$ | Antibiotic concentration for kill rate at half of its max ($\text{KC}_{50}$) | (3.7) | $\mu\text{g}/\text{ml}$ | (60) |
| $\xi$ | Density-dependent maximum increase in $\text{KC}_{50}$ | 10 | $\mu\text{g}/\text{ml}$ | (69) |
| $\gamma_k$ | Density for $\text{KC}_{50}$ at half of its maximum | $5 \times 10^6$ | CFU/ml | (69) |
| $\delta_B$ | Basal mortality rate | 0.05 | $\text{h}^{-1}$ | |
| $A_0$ | Peak antibiotic concentration in sputum | 0.05-100 | $\mu\text{g}/\text{ml}$ | (70) |
| $\delta_a$ | Antibiotic decay rate | 0.1 | $\text{h}^{-1}$ | (50) |
| $\lambda$ | Phage production rate | 1 | $\text{h}^{-1}$ | (13) |
| $\delta_V$ | Phage decay rate | 0.1 | $\text{h}^{-1}$ | (71) |

$a$ . see Supp. Material

### Analysis

All the analysis was performed using MATLAB. Growth rates were obtained directly from the equations above. We calculated the minimum inhibitory concentration (MIC) by setting Equation 5 to zero and solving for the antibiotic concentration. For simulations, we used the differential equation solver *ode45* to compute the bacterial density of Pf+ and Pf- strains over a minimum of 5 days, for varying number of doses per day and antibiotic concentration per dose. We then used the log-ratio of the median bacterial density for each strain in the last day of each simulation as a metric for comparison across different antibiotic regimens. We assumed that infection was cleared when bacteria’s density dropped below the quasi-extinction threshold of 1 CFU/ml. If the density of both strains decreased below this threshold, we then set their log-ratio to zero as well.

### Data availability

All code is available on Github at jpourtois/phage-antibiotics.

## Results

### Fitness of Pf+ and Pf- strains in the absence of antibiotic treatment

#### In the absence of antibiotics, Pf- reaches higher densities than Pf+. It also drives Pf+ to extinction when they coexist in the same infection sites

The metabolic cost of phage production *θ* leads to a reduction in Pf+ per-capita growth rate and, ultimately, in a lower long-term equilibrium for Pf+ strains relative to Pf- strains (Figure 2A). When in isolation (indirect competition), both strains grow to densities above 10^7^ CFU/ml, consistent with the number of genomic copies observed in CF lungs (7, 35). Pf+ and Pf- strains eventually reach their respective carrying capacity in independent infection sites: the carrying capacity of Pf- is equal to *K* · (1 – *δ_B_*/*r_max_*) and exceeds Pf+’s carrying capacity—equal to *K* · (1 – *δ_B_*/(*r_max_*(1 – *θ*)))—due to the extra energetic cost *θ* of phage production for Pf+ strains.

**Figure 2.**
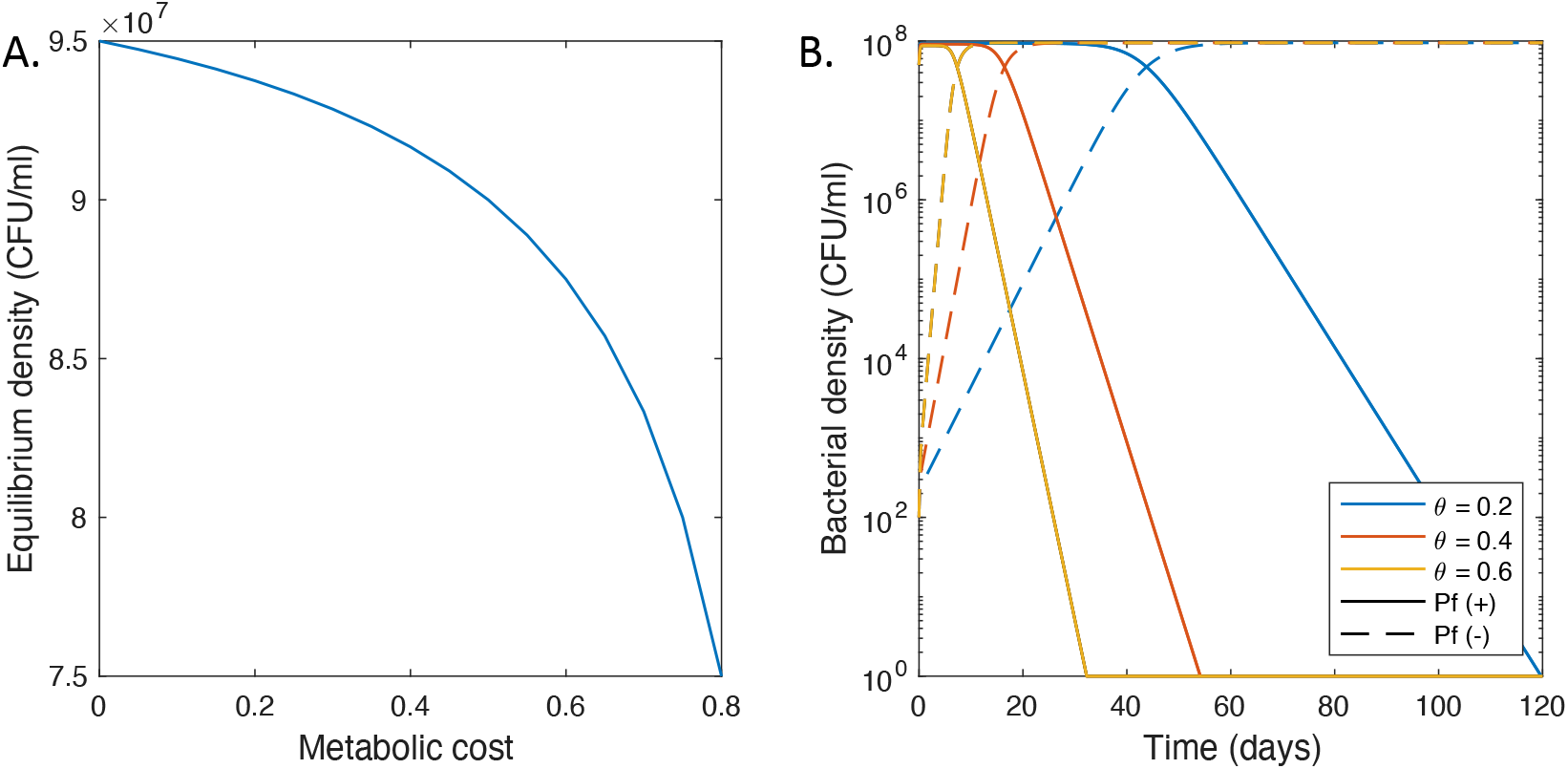
Growth dynamics of Pf- and Pf+ strains in the absence of antibiotics. **A.** Bacterial density at equilibrium versus metabolic cost. **B.** A plot of the predicted bacterial densities of Pf+ (solid curves) and Pf- (dashed curves) bacteria versus time following the introduction of 100 Pf- individuals into a Pf+ colony at equilibrium. We consider three different metabolic costs for Pf+ strains.

If a small population (10^2^ CFU/mL) of Pf- bacteria is introduced and directly compete with an established Pf+ colony, assuming *θ* > 0, the density of the Pf- strain increases over the course of multiple days until it reaches a density close to that of the Pf+ (Figure 2B). The density of the Pf+ strain then decreases before eventually dropping below the extinction threshold of 1 CFU/ml. The time required for Pf- strains to outcompete Pf+ bacteria decreases with increasing metabolic cost of phage production. For a metabolic cost of 0.6 (corresponding to a sixty percent reduction in growth rate), it takes 10 days for the Pf- strain to exceed Pf+ bacteria’s density and 32 days to drive the Pf+ strain extinct. Time to quasi-extinction increases to 54 and 119 days for a metabolic cost of 0.4 and 0.2, respectively.

### Antibiotic treatment under direct competition between Pf+ and Pf- strains

#### Pf+ cannot outcompete Pf- if both benefit from the sequestration of antibiotics by phages

When competing directly for space and resources, Pf+ and Pf- bacteria strains experience the same phage and antibiotics concentrations in their environment. In this section we assess antibiotic effect on both strains living in direct competition and we explore two antibiotic scenarios, namely: when the antibiotic effect is not dependent on the replication rate of the target bacteria (*ϵ_r_* = 0, eq. 7), and when it is dependent on replication rate (*ϵ_r_* = 1).

When the antibiotic effect is not dependent on replication rate (*ϵ_r_* = 0), the population growth rate of Pf- strains remains higher than the growth rate of Pf+ strains across the range of antibiotic concentrations (Figure 3A). In this scenario, the Pf+ strain bears the energetic cost of phage production while both strains benefit from the lower effective antibiotic concentrations due to phage sequestration of antibiotic molecules. As a consequence, Pf- outcompetes Pf+ at any antibiotic concentration that does not extirpate both strains.

**Figure 3.**
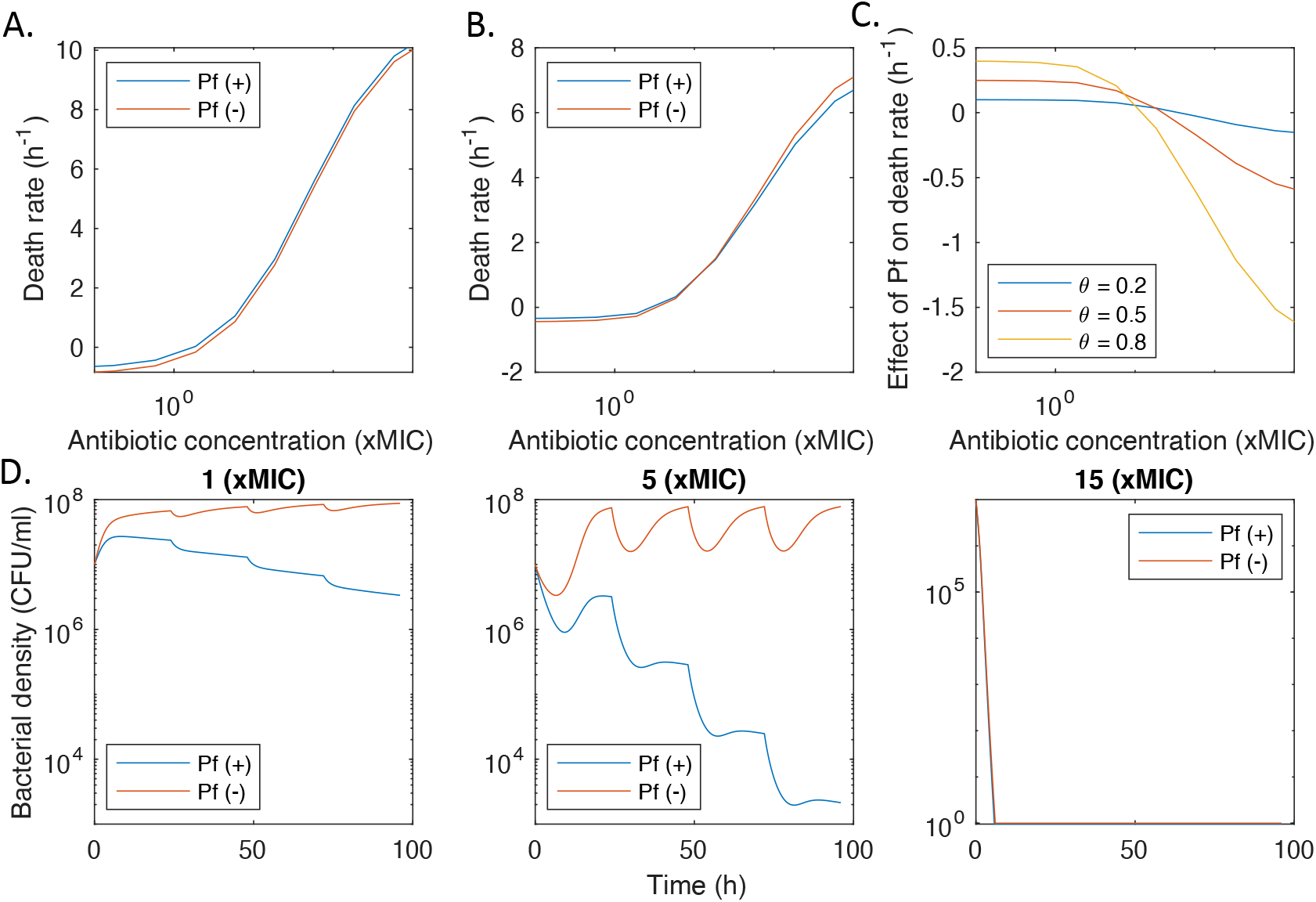
Growth dynamics of phage-negative and phage-positive strains in direct competition during antibiotic treatment. **A.** Plot of the net death rate of Pf+ and Pf- bacteria when the killing rate of the antibiotic is not dependent on replication rate. **B.** Plot of the net death rate of Pf+ and Pf- bacteria when the killing rate of the antibiotic is dependent on replication rate. **C.** Effect of phage production on net death rate for different costs of phage production (θ). This effect is the difference between the growth of Pf+ and Pf- strains. **D.** Bacterial density over time when one dose (of the indicated concentration) of a replication-dependent antibiotic is given per day.

We also investigated whether competitive exclusion of Pf+ strains coexisting with Pf- strains in the same infection sites can be prevented when antibiotics target specifically bacteria’s replication pathways (*ϵ_r_* = 1). In such a case, the benefit of phages’ protective biofilm is still equally shared between Pf+ and Pf- strains. However, the energetic cost *θ* of phage production (eq. 2), sustained by Pf+ strains only, might be partially compensated by the strain-specific reduced sensitivity to antibiotics due to Pf+ lower reproductive rate (eq. 7). Simulations showed that, up to 10 xMIC, Pf- continues to have a lower death rate than the Pf+ strain. However, Pf+’s lower replication rate leads to a relative advantage compared to Pf- strains at higher antibiotic concentrations (Figure 3B). The difference in death rate between Pf- and Pf+ at these elevated antibiotic concentrations increases as the metabolic cost of phage production, *θ*, increases (Figure 3C). The reduced reproductive rate of Pf+ bacteria caused by phage production thus provides an additional mechanism for tolerance against antibiotics with replication-related targets. However, this additional benefit is insufficient to compensate for the reduction in replication rate and, ultimately, to flip the outcome of competition between Pf+ and Pf-. Therefore, for a realistic range of model parameters analyzed in this work, under direct competition Pf- systematically outcompetes Pf+ also in the case of replication-targeting antibiotics.

As case studies, we compared the population dynamics of both strains for three different peak concentrations of antibiotics, administered once a day. At both 1 and 5 xMIC of antibiotics per day, the Pf- strain is able to maintain a high density, while the density of Pf+ shows decreases and recovery in response to the treatment with an overall decrease in density over time (Figure 3D). This would eventually result in the extinction of the Pf+ strain if the treatment were extended. At 15 xMIC per day, both strains rapidly decline to zero following the first dosage.

### Antibiotic treatment in the case of non-overlapping infection sites (indirect competition)

#### The effect of phages on Pf+ fitness depends on the sequestration constant, the decay rate of the antibiotic, as well as its dependence on bacterial replication

In the case of non-overlapping bacterial strains (indirect competition), phages are absent in Pf- strain environments and, thus, there is no change in the effective antibiotic concentration experienced by Pf- strains. In Pf+ infection sites, on the contrary, phages can sequester some amount of the antibiotics, essentially lowering the local concentration of antibiotics acting on Pf+ bacteria. However, in Pf+ environments the phage biofilm may exert different protection levels depending upon the specific type of antibiotic (7). First, we investigated how different values of the antibiotic sequestration factor *ϕ* (eq. 8) may affect the dynamics of bacterial infection. In order for antibiotic sequestration by phages to have a measurable impact on the growth rate of the Pf+ strain, our simulations show that the sequestration factor must exceed 10^5^ antibiotic molecules per phage (Figure 4A and S3, Supplementary text). A sequestration factor of 10^7^ led to a larger decrease in killing rate, especially at concentrations around 10 xMIC (Figure 4A). The degree of dependence of the killing rate on the replication rate *ϵ_r_* also affected the difference in growth rate between the Pf+ and the Pf- strains (Figure 4B). Pf+ strains showed a slower death rates than Pf- above 100xMIC only for antibiotics that target replication, consistent with our previous findings (Figure 3A and 3B). Finally, the density of Pf+ relative to Pf- increased for lower antibiotic decay rate (Figure 4C). Antibiotics with a low decay rate favor Pf+ strains because the antibiotic concentration is maintained for a longer period, which can lead to extinction of the Pf- strain. This is valid up to a certain concentration of antibiotics that is lethal to both strains.

**Figure 4.**
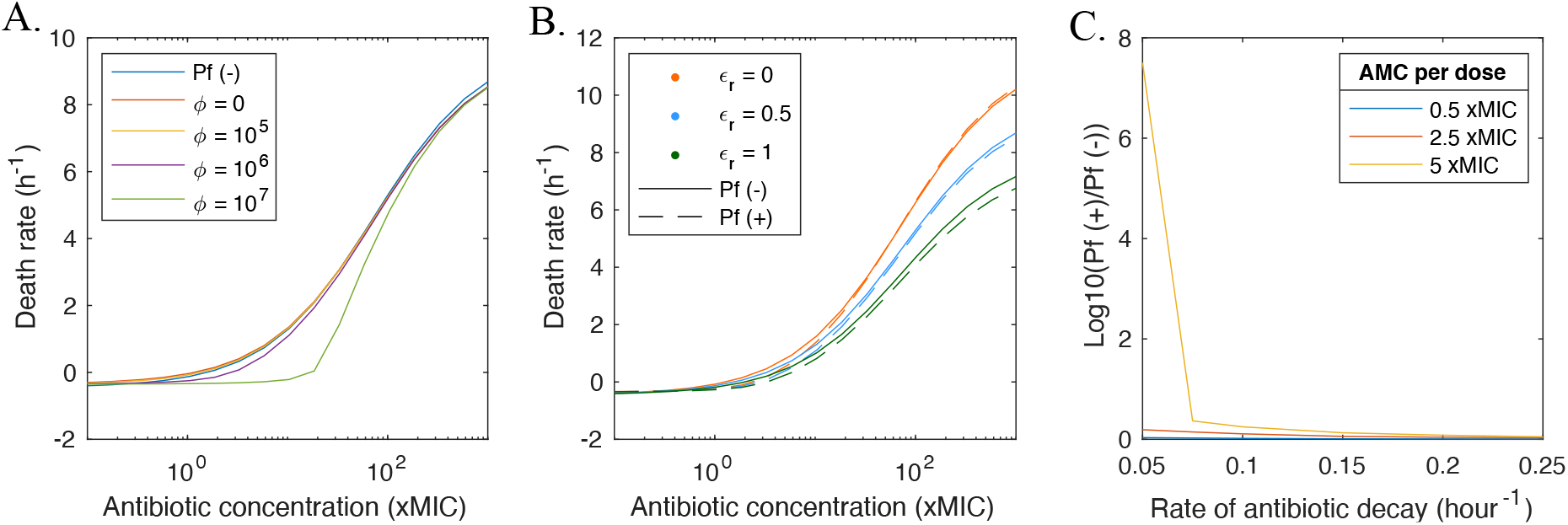
Effect of antibiotic properties on growth dynamics of Pf+ and Pf- strains in indirect competition. **A.** Plot of the net death rate of Pf+ and Pf- strains at different levels of antibiotic sequestration (ϕ). **B.** Growth rate of Pf+ and Pf- bacteria for antibiotics with low, intermediate and high levels of dependence on growth rate for efficacy (ε_r_). **C.** Comparison of the bacterial density of phage-negative and phage-positive strains for antibiotic regimens with different rates of antibiotic decay and antibiotic concentrations per dose (one dose per day). This comparison is expressed as the log of the ratio of the median over the last day of Pf+ to Pf- bacterial density.

#### In indirect competition, Pf+ bacteria outperform Pf- bacteria at concentrations of antibiotics close to or slightly higher than MIC for replication-dependent antibiotics

Next, we evaluated the effects of different concentrations of a replication-dependent antibiotic (*ϵ_r_* = 0.5, eq. 7) on Pf+ and Pf- bacterial strains in indirect competition. We consider regimens with different concentrations per dose and varying number of doses per day. In these conditions, Pf- bacteria have a higher growth rate at concentrations much lower than MIC but became less competitive than Pf+ bacteria at concentrations close to MIC or higher (Figure 5A). This resulted in Pf+ reaching higher densities than Pf- at higher concentrations of antibiotics per day and administered in fewer doses (Figure 5B). For fixed concentrations per dose, the relative fitness of Pf+ increased with the number of doses for 0.5 xMIC per dose (Figure 5C). At the intermediate concentration of 2.5 xMIC, the relative fitness of Pf+ increased first, before decreasing as the number of doses per day increased. Finally, both strains went extinct when a dose of 5 xMIC was used more than once a day. In other words, the fitness benefit of producing phages is greatest during treatment regimens with the highest exposure of antibiotics (i.e., highest concentration per dose, number of doses, and total duration of treatment), up to a point where phages cannot sequester enough antibiotics to prevent first a decrease in growth rate and then extinction.

**Figure 5.**
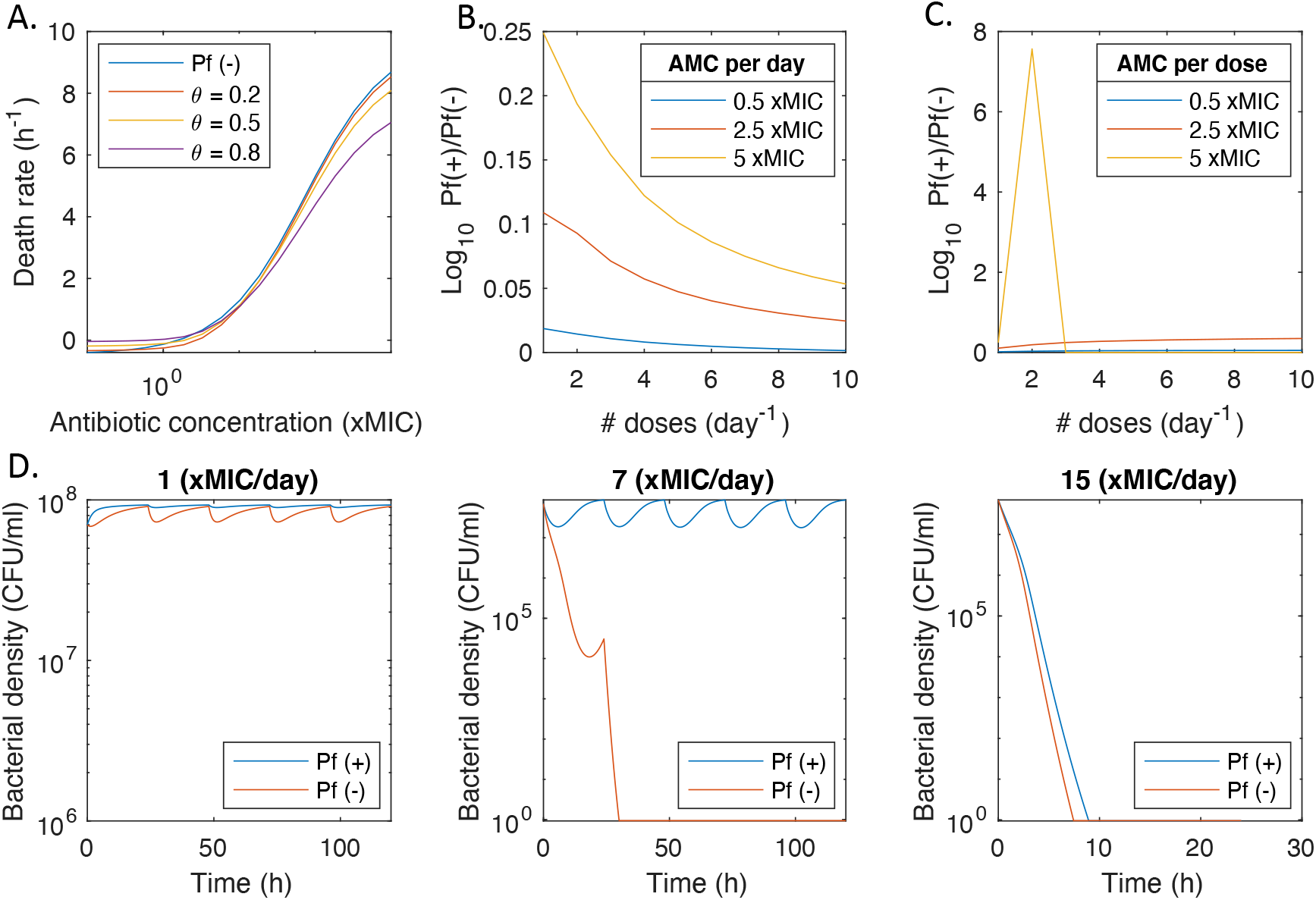
Growth dynamics of Pf+ and Pf- strains in indirect competition during antibiotic treatment. **A.** The net death rate of Pf+ and Pf- bacteria for different metabolic costs (θ) and antibiotic concentrations. **B.** Comparison of the bacterial density of Pf+ and Pf- strains for antibiotic regimens with different number of doses per day and total antibiotic concentrations per day. This comparison is expressed as the log of the ratio of the median over the last day of Pf+ and Pf- bacterial density. The ratio is set to 0 if both strains go extinct. **C.** Comparison of the bacterial density of Pf+ and Pf- strains for antibiotic regimens with different number of doses per day and antibiotic concentrations per dose. This comparison is expressed as the log of the ratio of the median of Pf+ and Pf- bacterial density over the last day. The ratio is set to 0 if both strains go extinct. **D.** Bacterial density over time when one dose of antibiotics is given per day, for three different doses.

We chose three different antibiotic concentrations to illustrate three possible competition outcomes when administered once a day (Figure 5D). At high antibiotic concentrations (15 xMIC), both strains go extinct. At an intermediate antibiotic concentration of 7 xMIC per day Pf+ bacteria maintained a density above 10^6^ CFU/ml, while Pf- went extinct over the course of five days. Finally, Pf+ and Pf- both survive the antibiotic treatment at a lower antibiotic concentration of 1 xMIC, with the density of Pf+ being higher, on average, than the density of the Pf- strain. It is possible for Pf- to over compete Pf+ for certain combinations of parameters (Figure S1). This occurs when the death rates associated with antibiotics is low (Figure S2). In these conditions, the benefits provided by phage sequestration are not large enough to justify the metabolic cost.

## Discussion

Although most bacteriophages have traditionally been thought to have only parasitic relationship with their bacterial hosts, the discovery of commensalistic and even mutualistic phage-bacteria relations suggests that a deeper understanding of phage biology is clinically important. With the discovery that filamentous phages lead to more resistant biofilms and protection against many antibiotics, we set out to use a mathematical approach to investigate how Pf phages impact the fitness of Pa in different antibiotic environments. Namely, we wanted to understand what factors may be contributing to the emergence and resilience of Pf+ Pa infections. These findings could also prove informative for other phage-producing bacteria such as *Escherichia coli* and *Klebsiella pneumonia*.

We found that in the absence of antibiotics, Pf- bacterial strains outcompeted Pf+ strains, as expected given the energetic cost of phage production, an outcome well known in ecology as the principle of competitive exclusion for species exploiting the same resources, or also Gause’s Law (53). This is consistent with the low Pf phage prevalence in pediatric patients with CF, many of whom have received relatively limited antibiotic treatments (7). In direct competition, assuming that Pf- strains cannot be infected by phages, Pf+ strains are unable to outcompete Pf- strains even under antibiotic treatment, because Pf+ strains sustain the energetic cost of phage production, but share the benefits evenly with Pf- strains.

Overall, our model suggests that Pf+ can outcompete Pf- only when Pf+ strains do not overlap in the same infection sites with Pf- and thus, while bearing the cost of phage production, they do not share the benefit with Pf- strains. A recent report demonstrated that Pf+ bacteria grown *in vitro* may encase themselves within bundles of phage, thus providing a shielding mechanism against antibiotics (54). It is unclear, however, if this behavior occurs in clinical settings or if Pf+ bacteria can protect Pf- bacteria against antibiotics within a mixed-strain infection. It is conceivable that Pf+ strains may be able to outcompete Pf- strains due to this highly localized phage presence. Our model simplifies these questions by using strict definitions of direct and indirect competition. Further studies are necessary to understand these co-culture biofilms, particularly in clinically relevant settings.

Interestingly, Pf+ strains are not able to dominate in all indirect competition scenarios. Intermediate concentrations of antibiotic are critical for Pf+ to significantly outcompete Pf-. These concentrations correspond to the regime at which antibiotic sequestration by phages lowers the effective antibiotic concentration thus lowering the death rate of Pf+ bacteria enough to compensate for the higher energetic cost of phage production. Under these circumstances, Pf+ density exceed that of Pf-. Below these antibiotic concentrations, the marginal benefit of a reduction in antibiotic effective concentration accrued thanks to the protective effect of the phage biofilm is unable to compensate for the sheer cost of phage production and thus Pf+ density remains lower than that of Pf- strain. At high antibiotic dose, the ability of phages to sequester antibiotic is overwhelmed and infections with both Pf+ and Pf- are eventually cleared. This is consistent with the clinical evidence that shows Pf+ strain infections correlated with chronic infections (7) that have been treated with antibiotics. Once a patient with CF has failed eradication, chronic inhaled anti-pseudomonal therapy is initiated with inhaled tobramycin, or alternatively aztreonam. While the sputum antibiotic levels achieved with inhaled antibiotics are reported as well above MIC (55, 56), it is plausible that some areas of infected lung tissue do not reach such high levels. CF is an obstructive pulmonary disease which is characterized by areas of heterogeneous ventilation (57, 58). There are likely distal areas of the lung with poor ventilation that do not see the same delivery of inhaled antibiotics, thus creating an environment with intermediate concentrations of antibiotics favoring the survival and dominance of Pf+ strains. The continuous use of antibiotics at sub-lethal concentrations may thus be partially responsible for driving infections towards a Pf+ dominated state. Our model would suggest that an ideal treatment for avoiding Pf+ infections would use antibiotics that cannot be sequestered by phages, and do not work through a replication-dependent mechanism.

In this model, we have focused on the effects of phages on biofilm-embedded bacterial strain fitness under antibiotic stress. However, we have not accounted for how the presence of phages may change the properties of the biofilm with respect to antibiotic treatment. For example, it is known that phage-rich *Pa* biofilms show liquid crystal structures but it is unclear how this organization and the associated changes in the chemical potential of the biofilm may contribute to antibiotic tolerance (33). Furthermore, Pf phages may also impact other critical properties of the infection site, such as the rheological properties of the infection environment, the adhesivity of the biofilm, and the transport of nutrients and antibiotics. Finally, we focus here on antibiotics delivered orally as part of medical treatment. However, antibiotic substances can also be secreted by bacteria to prevent interspecies competition (59). Our results could thus inform our understanding of bacterial competition in a wide range of context, as well as in the clinical setting we focus on in this work.

## Conclusion

Filamentous phages have increasingly been recognized as important players in the development of chronic infections through their effect on antibiotics, immunity and biofilm formation. In particular, Pf+ infections in CF patients make up a larger fraction of chronic infections and lead to worse lung functions. Our simulations showed that the fitness of Pf+ strains is highest when antibiotic sequestration by phages is high but localized, and when a constant intermediate concentration of antibiotics is maintained in the lungs. This suggests that the frequent antibiotic treatments associated with CF could contribute to the increased prevalence of Pf+ strains in CF patients. Not all antibiotics have the same properties, however, and our model suggests that high sequestration, replication-dependent mechanisms and low decay rates particularly favor Pf+ strains. Of note, phages have additional effects on bacteria and the human body that are still poorly understood and could not be captured in our model. A better understanding of the ecology of phages in the human body will be an essential step for developing more successful treatment strategies of bacterial infections.

## Acknowledgments

JP acknowledges support from the Stanford Graduate Fellowship. PLB is supported by grants from the Falk Medical Research Trust, R01 AI138981-01, R01 HL148184-01, grants from Stanford Bio-X, SPARK, and the CFF.

We would also like to acknowledge Jonas D. van Belleghem for many helpful discussions and Ajai A. Dandekar for his comments and suggestions during the writing period.

## Author Contributions

JDP and PLB conceived of the presented idea. JDP designed the model and performed the analysis. GADL contributed to revise, update and fine tune the modeling section. All authors performed research with regard to model assumptions and parameter values. All authors contributed to the graphics and writing of the manuscript.

